# QTL Mapping and integration as well as candidate gene prediction for branch number in soybean

**DOI:** 10.1101/2019.12.27.889287

**Authors:** Yuhua Yang, Yang Lei, Zhiyuan Bai, Yichao Wei, Ruijun Zhang

## Abstract

Branch number is an important factor that affects crop plant architecture and yield in soybean. With the aim of elucidating the genetic basis of branch number, we identified 10 consensus quantitative trait loci (QTLs) through preliminary mapping, which were on chromosome A1, B2, C1, C2, D1a, D1b, F, L and N, explained 0.3-33.3% of the phenotypic variance. Of these, three QTLs were identical to previously identified ones, whereas the other seven were novel. In addition, one major QTL-*qBN.C2* (R^2^=33.3%) was detected in all three environments and another new major QTL-*qBN.N* (R^2^=19.6%) was detected in two environments (Taiyuan 2017 and Taiyuan 2018), but only in Taiyuan. Thus, the QTL × environment interaction analysis confirmed that QTL-*qBN.N* was strongly affected by the environment. We compared the physical positions of the QTL intervals of the candidate genes potentially involved in branching development, and five orthologous genes were ultimately selected and related to the establishment of axillae meristem organization and lateral organs, *qBN.A1* (SoyZH13_05G177000.m1), *qBN.C2* (SoyZH13_06G176500.m1, SoyZH13_06G185600.m1), and *qBN.D1b-1* (SoyZH13_02G035400.m1, SoyZH13_02G070000.m3). The results of our study reveal a complex and relatively complete genetic architecture and can serve as a basis for the positional gene cloning of branch number in soybean.

## Introduction

Plant architecture is an extremely important role in plant yield, which can affect light distribution in the canopy and photosynthesis. Modification of crop plant architecture has been used to improve plant fitness and agricultural performance, and these modifications are achieved through genetics and breeding [1–3]. Branching is a major factor that affects plant architecture, together with plant height, main stem, leaf, pod and others [2]. The number and distribution of branches determine the canopy architecture, which influences light interception as well as lodging resistance and ultimately seed yield [4]. Branch number is strongly dependent on aspects related to the cultural practices and growth environment [5].

Branch number is also the most complex trait and is very sensitive to the environmental conditions. Because the environmental effects on branch number are so prominent, there have been few studies on the genetics of this trait, although varietal differences in branch number have been observed in the adaptation to row spacing in accordance with its modest heritability observed in most studies [6, 7]. Two dominant alleles at independent loci were found to be associated with a high-branch number phenotype [8] but the loci and other genetic factors that determine branch number in soybean remain unknown. However, branch number is difficult to improve efficiently by traditional breeding methods and the genetic and especially the molecular mechanism involved in branch number in soybean are poorly understood.

Previous studies have shown that branch number is controlled by multiple quantitative trait loci (QTLs), and almost 93 branch number QTLs have been identified from nearly 18 linkage mapping populations [9–20]. Interestingly, a few QTLs on the C2 chromosome were found to be concentrated in a region (103-121cM) derived from different parents and environments [4, 21–25]. It is indicated that this region on the C2 chromosome is a hotspot for branch number in soybean. However, only a few of these QTLs have been repeatedly detected, in accordance with the modest heritability of branch number. In addition, almost all these branch number QTLs were found to have a moderate effect. Therefore, these data suggest that the existence of varietal differences in branch number among soybean cultivars and it is difficult to narrow down these QTLs and identify the underlying candidate genes.

The objectives of the present study were to (a) identify the genetic mechanism that controls branch number in soybean by performing quantitative trait locus (QTL) analysis using F_2_ population derived from two soybean lines, C025 and JD18, that have showed consistent significant differences in branch number in different environments, to (b) dissect the relationship between major QTLs and the environment, to (c) integrate QTLs associated with branch number in soybean using meta-analysis and to (d) predict the potential candidate genes for further fine-mapping and mechanism studies.

## Materials and methods

### Plant materials, field experiments and trait investigation

The F_2_ population included 109 individuals, and was derived from two soybean lines, C025 (high branch number) and JD18 (low branch number). The F_2_ individuals were planted in Hainan from Nov. 2016 to Feb. 2017 (code Hainan 2016), the F_2:3_ and F_2:4_ lines were planted in Taiyuan from May 2017 to Oct. 2017 (code Taiyuan 2017) and May 2018 to Oct. 2018 (code Taiyuan 2018). The F_2_, F_2:3_ and F_2:4_ populations and two parents were arranged in a randomized complete block design with two replications. Each block contained two rows with a space of 50 cm between rows and 13.5 cm between individual plants. The seeds were sown by hand, and the field management followed standard agriculture practices. In each block, 10 representative individuals from the two rows were harvested by hand at maturity. The branch number was measured based on a previously described method [4].

### SSR molecular marker analysis and linkage map construction

The SSR markers from the SoyBase database (https://soybase.org/) were used for polymorphism screening between the two parents and used to genotype the F_2_ individuals. Leaf tissue was collected from seedlings of the parents and F_2_ populations. Genomic DNA was extracted according to the CTAB method [26]. The PCR, electrophoresis, and silver staining procedures were performed as described previously [27].

The genetic linkage map was constructed using the software JoinMap 4.1 (https://www.kyazma.nl/index.php/JoinMap/) with a threshold for goodness-of-fit ⩽ 5, recombination frequency of < 0.4 and minimum logarithm of odds (LOD) score of 2.0. The genetic distances were measured based on the Kosambi function. To avoid the potential errors, double-crossover events were checked.

### QTL mapping and meta-analysis

The composite interval mapping (CIM) method was used for QTL mapping, and was incorporated into WinQTLCart v2.5 software (http://statgen.ncsu.edu/qtlcart/WQTLCart.htm). The walking speed, number of control markers, window size and regression method were set to 1cM, 5, 10cM and forward regression, respectively. The LOD threshold was determined by 1000 repetitions of permutation analysis [28]. The LOD value corresponding to P=0.05 (3.0-4.6) was used to detect significant QTLs. Moreover, a lower LOD value corresponding to P=0.10 (2.6–4.1) was employed to identify suggestive QTLs with small effects.

Meta-analysis was performed to estimate the number and position of the true QTLs underlying the QTLs of the same or related traits, which were repeatedly detected in the different environments and/or populations [29]. The computation was conducted using BioMercator 2.1 software [30]. Each consensus QTL was designated with the initials “cq”, followed by the name of the trait abbreviation and linkage group.

### Statistical analysis

The broad-sense heritability was calculated as *h*^2^= *σ*2 g/(*σ*2 g + *σ*2 ge / *n* + *σ*2 e / *nr*), as described previously, where *σ*2 g, *σ*2 ge, and *σ*2 e are the variance of genotype, genotype×environment, and error, respectively, and n and r are the number of environments and replications, respectively. The values of *σ*2 g, *σ*2 ge, and σ2 e were estimated using the SAS ANOVA procedure.

## Results

### Phenotypic variation of the parents and F_2_ population

The two parents, C025 and JD18, showed extremely significant differences in branch number in all three investigated environments. The branch number of C025 was 4.1 ± 0.83, which was approximately twice the branch number of JD18 (1.6± 0.79) (Table 1). The branch number of the F_2_ population showed more or less normal distributions in all three investigated environments (Fig 1), indicating a quantitative inheritance suitable for QTL identification. In addition, the branch number of the F_2_ population exhibited transgressive segregation but to a small degree, indicating that the favorable alleles were mainly distributed in one of the two parents.

**Fig 1.**
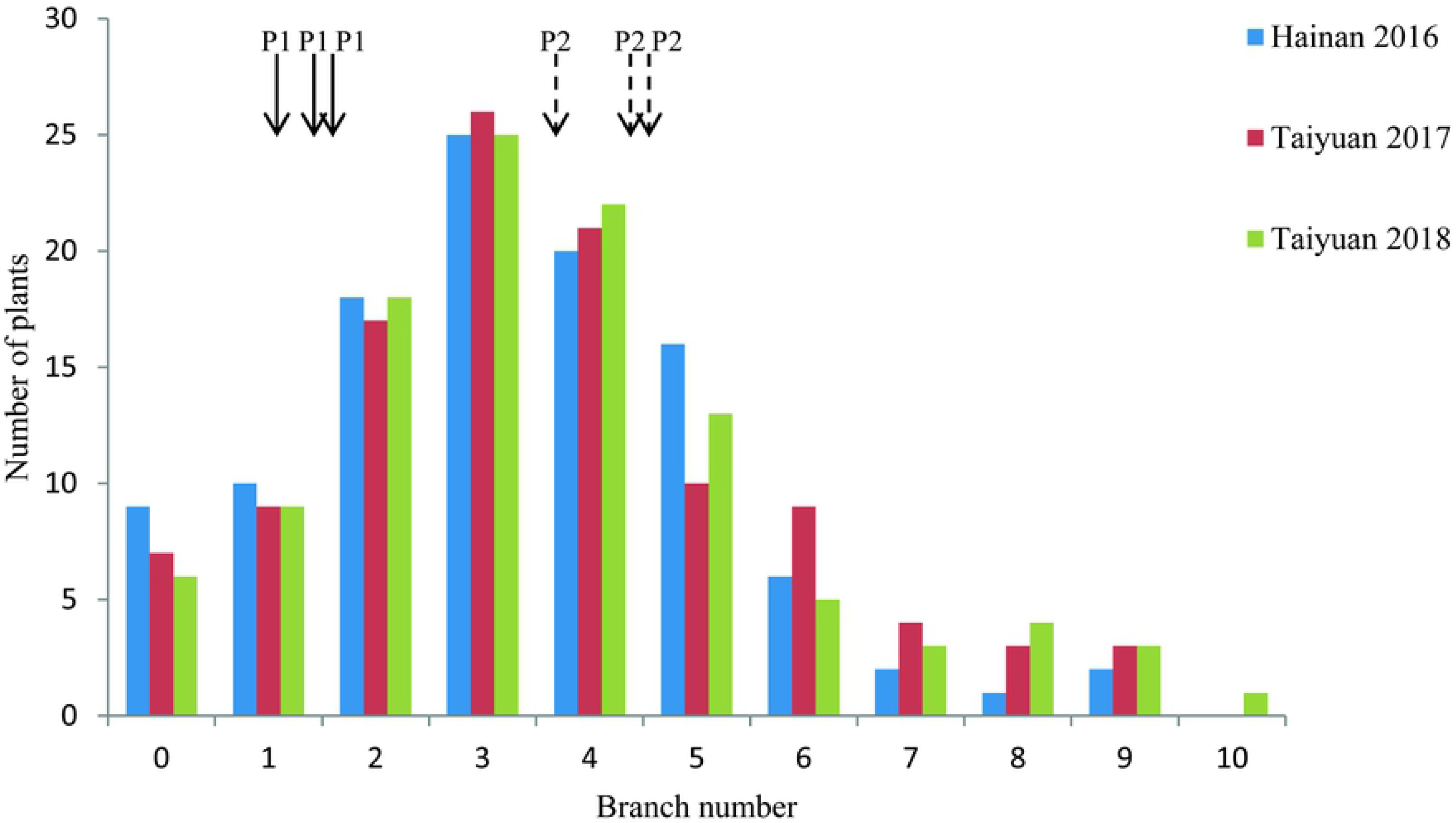
Frequency distribution of branch number for the F_2_ population in three environments. The horizontal axis represents the trait value of branch number. The vertical axis represents the number of individuals. P1 and P2 indicate H18 and C025, respectively.

**Table 1.** Phenotypic variation of branch number for the parents and F_2_ population in three investigated environments

| Environment code | Parents |  |  | F<sub>2</sub> population |  |  |
| --- | --- | --- | --- | --- | --- | --- |
|  | C025 | H18 | *P*<sub>t-test</sub> | Min | Max | Mean |
| Hainan 2016 | 3.6±0.29 | 1.2±0.81 | 3.8E-11 | 0.0 | 9.0 | 3.2±0.69 |
| Taiyuan 2017 | 4.4±0.92 | 1.8±0.27 | 2.9E-12 | 0.0 | 9.0 | 3.5±0.78 |
| Taiyuan 2018 | 4.4±0.47 | 1.7±0.66 | 1.5E-18 | 0.0 | 10.0 | 3.5±0.93 |
| Mean | 4.1±0.83 | 1.5±0.79 | 3.3E-36 | 0.0 | 9.3 | 3.4±1.09 |

The analysis of variance revealed that the genotypic, environmental and genotype × environment effects were all extremely significant for branch number (Table 2). The broad-sense heritability (*h^2^*) of branch number in the F_2_ population was 48%.

**Table 2.** ANOVA analysis and heritability estimation for branch number

| Source | Degree of freedom | Variance | Mean square | F value | P value | $\sigma^2$ | $h^2$ |
| --- | --- | --- | --- | --- | --- | --- | --- |
| Genotype | 108 | 2115 | 20 | 27.1 | <.0001 | 1.6 | 0.48 |
| Environment | 2 | 3931 | 1966 | 16.2 | <.0001 | 9.0 |  |
| Genotype by environment interaction | 216 | 2193 | 10 | 26.6 | <.0001 | 3.8 |  |
| Error | 326 | 824 | 3 |  |  | 2.5 |  |

### Linkage mapping of the QTLs for branch number in soybean

A common framework of the genetic linkage map containing 1015 SSR makers was constructed, which covered a total of 2282.9 cM of the soybean genome and had an average distance of 2.4 cM between adjacent makers (https://soybase.org/). A total of 13 branch number QTLs were detected and explained 0.3-38.3% of the phenotypic variance (S1 Table). Ten QTLs were integrated by the meta-analysis, and these were distributed on nine (A1, B2, C1, C2, D1a, D1b, F, L and N) of the 20 chromosomes. Of these, two repeatable consensus QTLs (*qBN.C2* and *qBN.N*) were distributed on the C2 and N chromosomes, *qBN.C2* was detected in all three environments (Hainan2016, Taiyuan 2017 and Taiyuan 2018), and *qBN.N* was detected in Taiyuan 2017 and Taiyuan 2018, and displayed a large effect (R^2^=33.3% and 19.8%), thus, they can be treated as major QTLs (Table 3). Of the eight other QTLs, *qBN.C1* was only detected in Hainan 2016, *qBN.D1b-1* was only detected in Taiyuan 2017, and the other six QTLs (*qBN.A1, qBN.B2, qBN.D1a, qBN.D1b-2, qBN.F* and *qBN.L*) were only detected in Taiyuan 2018.

**Table 3.** Detected QTLs for branch number of soybean from different environments

| QTL name | Chromosome | LOD value | *R*<sup>2</sup> (%) | Position | Confidence interval | Additive effect | Experiments code |
| --- | --- | --- | --- | --- | --- | --- | --- |
| *qBN.A1* | A1 | 6.5 | 10.3 | 79.9 | 76.3-84.1 | -0.60 | Taiyuan 2018 |
| *qBN.B2* | B2 | 2.6 | 5.2 | 39.9 | 38.4-42.5 | -3.15 | Taiyuan 2018 |
| *qBN.C1* | C1 | 3.7 | 2.9 | 1.0 | 0.1-4.0 | -0.22 | Hainan 2016 |
| *qBN.C2* | C2 | 4.7 | 33.3 | 105.9 | 98.7-110.3 | -1.27 | Hainan 2016,Taiyuan 2017,Taiyuan 2018 |
| *qBN.D1a* | D1a | 3.3 | 5.3 | 44.8 | 41.8-48.5 | -2.78 | Taiyuan 2018 |
| *qBN.D1b-1* | D1b | 2.5 | 8.9 | 18.0 | 1.9-39.9 | 2.02 | Taiyuan 2017 |
| *qBN.D1b-2* | D1b | 3.3 | 4.6 | 75.8 | 64.0-89.8 | -2.43 | Taiyuan 2018 |
| *qBN.F* | F | 2.7 | 11.6 | 10.0 | 0-18.0 | -1.87 | Taiyuan 2018 |
| *qBN.L* | L | 5.3 | 0.3 | 22.1 | 20.1-25.0 | -1.58 | Taiyuan 2018 |
| *qBN.N* | N | 2.9 | 19.8 | 84.6 | 76.5-87.6 | -2.46 | Taiyuan 2017,Taiyuan 2018 |

Interestingly, one major QTL-*qBN.N* was a novel branch number QTL in soybean, and was detected only in Taiyuan. Thus, to determine the relationship between QTL-*qBN.N* and the environment, a QTL × environment interaction analysis was performed (Fig 2). The results showed that the branch numbers of QQ and Qq-genotype in Hainan were lower than those in Taiyuan, but the branch number of the qq-genotype was higher, thus, these results indicated that QTL-*qBN.N* was strongly affected by the environment.

**Fig 2.**
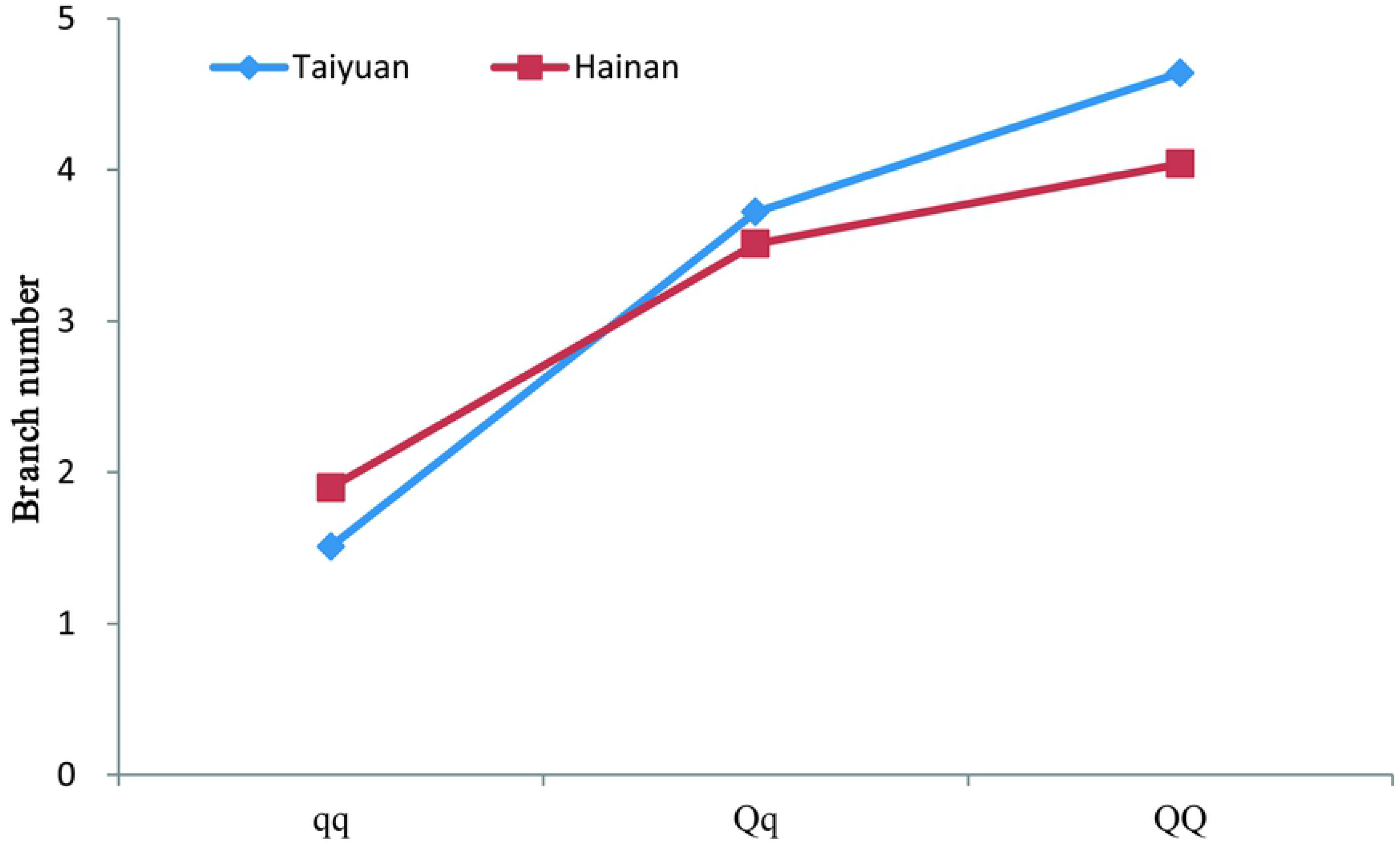
Interaction between *qBN.N* and the environment. The horizontal axis represents the different genotypes. The vertical axis represents the trait value of branch number.

### Meta-analysis of the QTLs for branch number

A total of 93 QTLs were identified in the reported QTL mapping (S2 Table), and 63 consensus QTLs (including current studies) were successfully anchored to a complete genetic map, which were subjected to meta-analysis (Table 4). These consensus QTLs were distributed on all of the 20 chromosomes. The R^2^ of these consensus QTLs ranged from 0.1 to 50.9%. About 80% QTLs showed small effects (R^2^ < 10%), and only four QTLs had large effects (R^2^ > 30%), which should be considered as major QTLs.

**Table 4.**
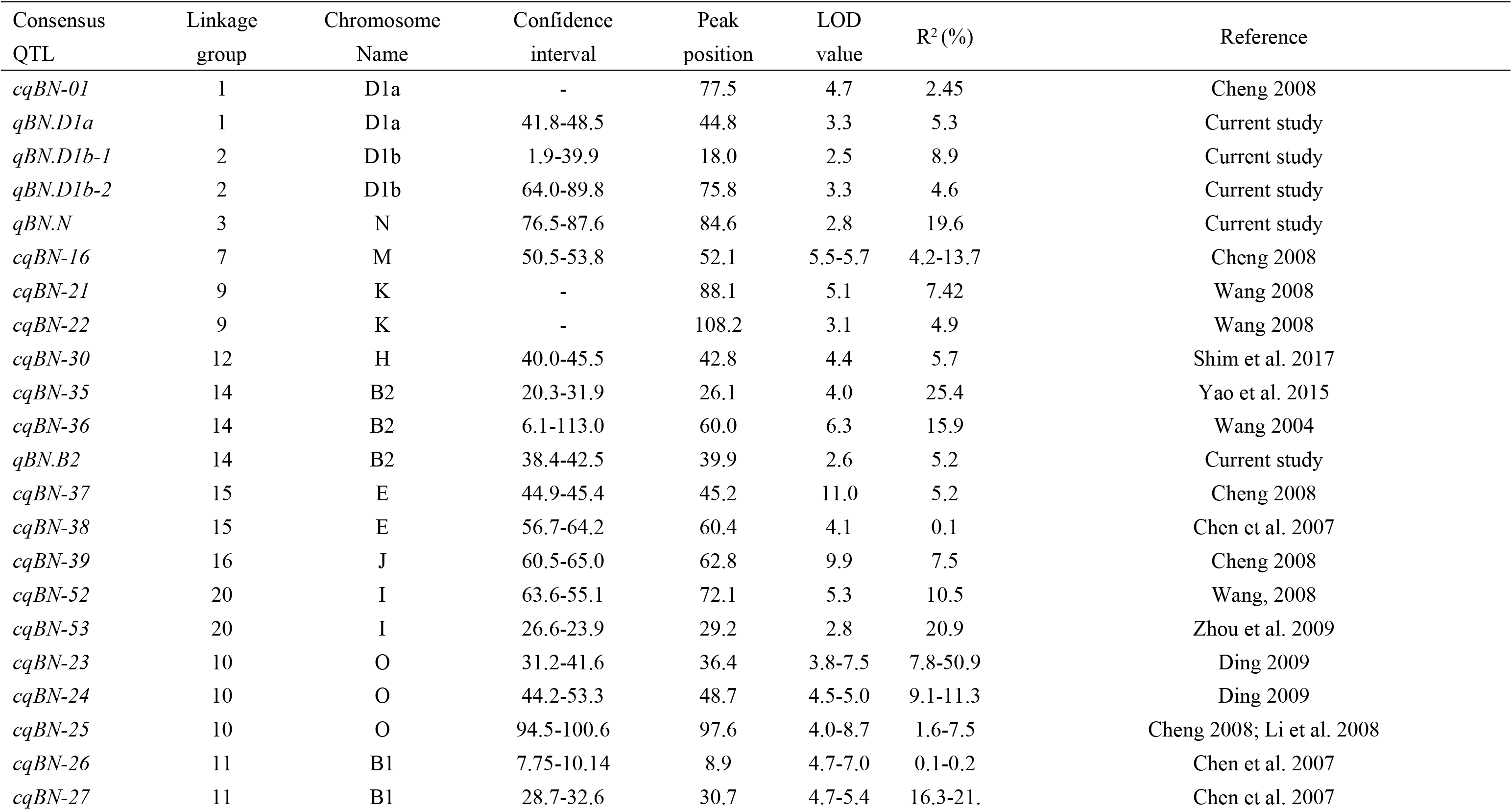
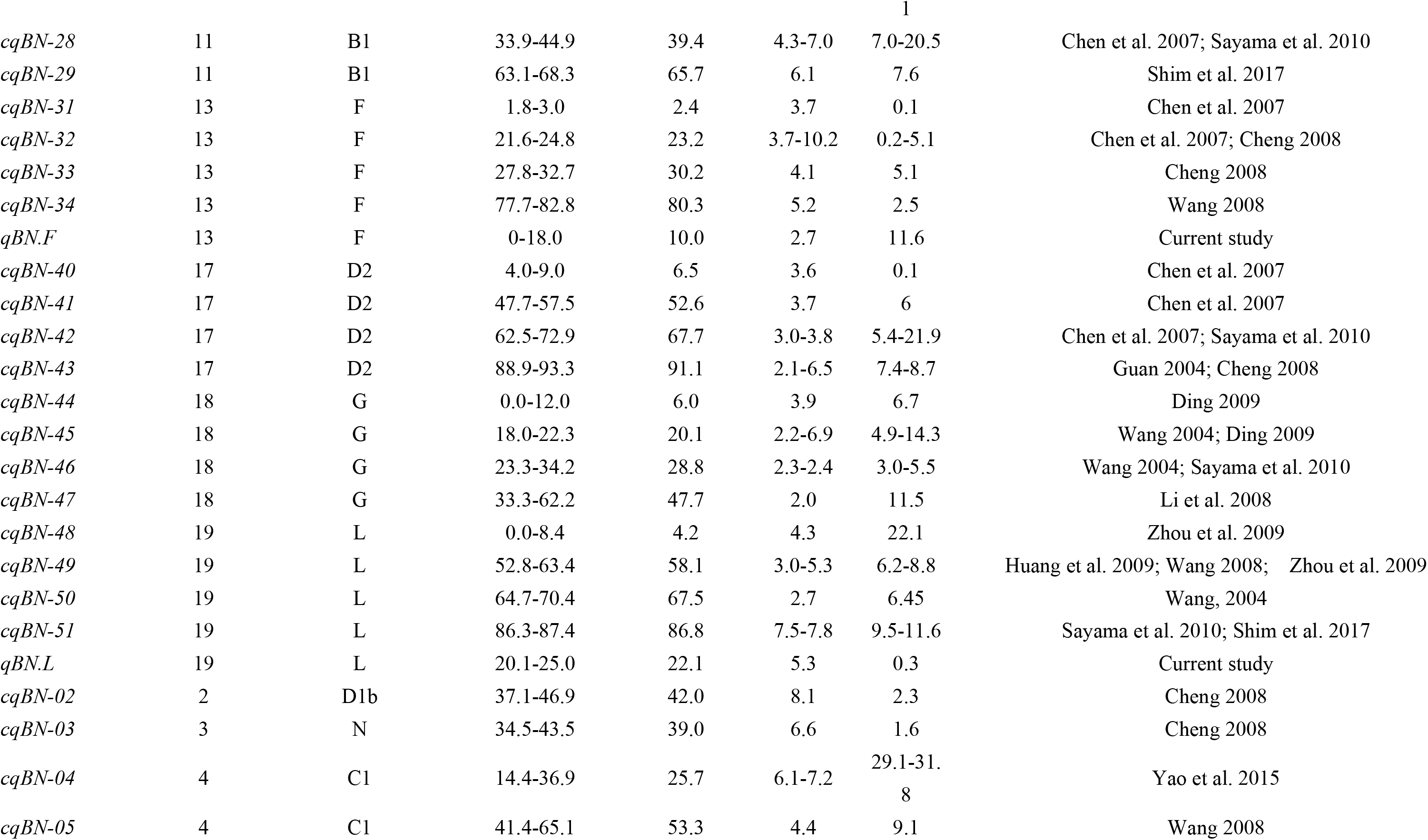
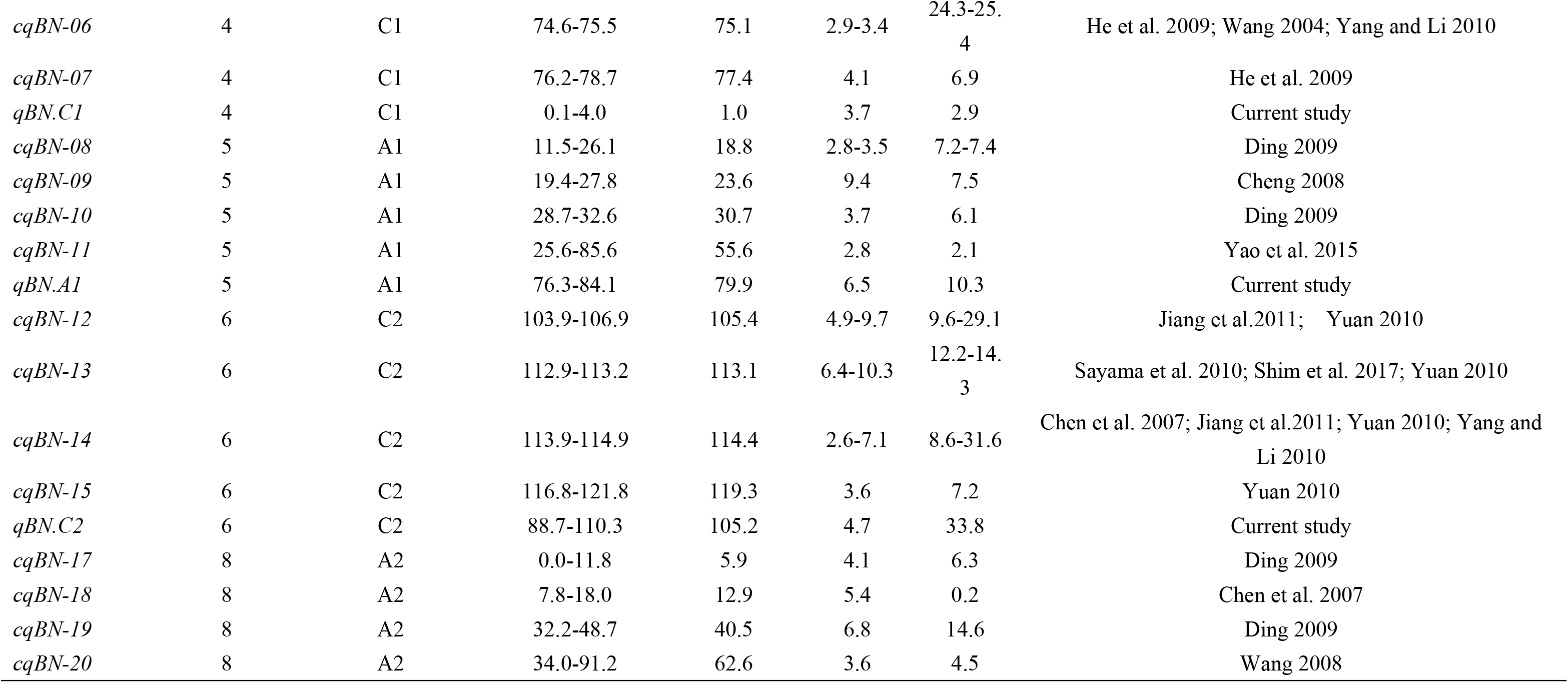
Common branch number QTLs detected in different studies

### Identification of candidate genes

To identify candidate genes for branch number in soybean, we extracted the gene sequences of 10 QTLs regions identified in the current study by blasting the bilateral sequence of SSR markers of QTLs to the Gmax_ZH13, which is the third generation sequencing of the soybean genome. After performing BLASTN searches against all of the genes in *A.thaliana,* we found five orthologous genes, which were involved in the establishment of axillae meristem organization and lateral organs and were possible candidate genes for branch number (Table 5). The five candidate genes resided in the regions of *qBN.A1* (*SoyZH13_05G177000.m1*), *qBN.C2* (*SoyZH13_06G176500.m1, SoyZH13_06G185600.m1*), and *qBN.D1b-1* (*SoyZH13_02G035400.m1, SoyZH13_02G070000.m3*) and have been shown to play an important role in regulating axillae meristem growth and development [24, 31–35]. For example, *SoyZH13_02G035400.m1* is homologous to Arabidopsis *cuc3* (*AT1G23380*), which encodes a NAG domain transcription factor and is a major regulator for axillary meristem initiation and separation of the meristem from the main stem [36–37]. SoyZH13_05G177000.m1 is homologous to Arabidopsis *knat6* (*AT3G18730*), one of KNOTTED-like homeobox (*KNOX*) genes, contributes to boundary establishment in the embryo via the *STM/CUC* pathway and matches the expression domain of the *CUC* genes in late embryogenesis [33]. The five genes should be considered as promising candidate genes that could be used for further study.

**Table 5.** Identification of candidate genes for branch number QTLs in soybean

| QTL name | Soybean Gene ID | Chromosome Name | Linkage group | Physical interval (Mb) | Homologous genes of Arabidopsis | Gene annotation |
| --- | --- | --- | --- | --- | --- | --- |
| *qBN.A1* | SoyZH13_05G177000.m1 | A1 | 5 | 39.329-39.339 | AT3G18730 | Encodes a protein with an important role in cell division control and plant morphogenesis and may play a role in genome maintenance. |
| *qBN.C2* | SoyZH13_06G176500.m1 | C2 | 6 | 16.108-16.113 | AT5G40280 | Encodes a beta subunit of farnesyl-trans-transferase, which is involved in meristem organization and ABA-mediated signal transduction pathway. |
| *qBN.C2* | SoyZH13_06G185600.m1 | C2 | 6 | 17.463-17.464 | AT1G76420 | Identified in an enhancer trap line; member of the NAC family of proteins. Expressed at the boundary between the shoot meristem and lateral organs and the polar nuclei in the embryo sac |
| *qBN.D1b-1* | SoyZH13_02G035400.m1 | D1b | 2 | 3.393-3.403 | AT1G23380 | Homeodomain transcription factor KNAT6, belonging to class I of the KN transcription factor family. |
| *qBN.D1b-1* | SoyZH13_02G070000.m3 | D1b | 2 | 6.407-6.413 | AT5G13570 | Encodes DCP2 with mRNA decapping activity. |

## Discussion

We analyzed the phenotypes of the F_2_ population in three environments and identified 10 branch number consensus QTLs. The results indicated that several regions, including those on chromosomes A1, B2, C1, C2, D1a, D1b, F, L and N are associated with the difference in branch number between C025 and JD18. Furthermore, the possible presence of multiple QTLs on chromosomes D1b suggests that branch number is controlled by multiple genes on the same chromosome. For such a trait, the development and use of mapping populations represents a potential powerful tool to reliably detect QTLs.

In previous linkage QTL mapping studies, approximately 93 QTLs were identified for branch number, which were distributed on all 19 chromosomes. Most of these QTLs showed relatively small effects, with only two major QTLs [17, 20], on the C1 and C2 chromosomes, respectively. Comparison of the regions of the QTLs detected in this study and meta-QTL for branch number reported by predecessor showed that *qBN.A1*, *qBN.C2* and *qBN.F* included meta-QTL, suggesting these three QTLs may have a robust effect on branch number in different genetic backgrounds if they are identical to these meta-QTL. These “repeatable” QTLs found across the current and previous studies should be potential targets for marker-assisted selection and map-based cloning. These results showed that branch number was controlled by a large number of loci, mostly with small effects, which strongly suggested the complexity of the genetic basis of branch number traits in soybean.

The effect of the QTL × environment interaction has also been addressed in several studies [38–40], including the result that the major QTL-*qBN.N* was detected only in Taiyuan. It has been suggested that a substantial proportion of QTL affecting a trait are active across different environments [41]. In previous studies, such *“*micro-real” QTL [42] or “ indicator QTL” [43], with positive or negative effects on traits were often ignored. However, such loci may be very useful to further increase yield under selection in different genetic backgrounds and environments [44].

We surveyed the presence of genes related to branch number reported by previous studies in detected QTLs candidate regions. Among the 10 QTL candidate regions, five candidate genes were predicted. Julier and Huyghe (1993) [45] suggested that the number of branches might be controlled by the number of axillary buds initiated. Thus, we found cell division-related genes such as those encoding a protein with an important role in cell division control and plant morphogenesis and transcription factors, including KNAT6, DCP2, ERA1, CUC3 and TSK. These might be candidate genes for branch number in soybean. To identify the causative genes, we are now performing high-resolution mapping. We expect that the use of these branch number-related genes may allow us to improve the soybean branch number system, resulting in high yields.

In the future, as data will be continually obtained through QTL mapping analyses [46, 47], it will be possible to integrate the data from this study into further meta-analyses using this approach to provide a better understanding of complex traits for crop improvement. These data will provide an important theoretical and applied resource for studies on soybean genetics and genomics.

## Conclusion

Total of 10 QTLs for branch number were detected in F2/F_2:3_/F_2:4_ populations using a consensus linkage map, two of which (*qBN.C2* and *qBN.N*) were validated in repeatable populations, and indicated that QTL-*qBN.N* was strongly affected by the environment. In addition, 5 candidate genes were ultimately selected and related to the establishment of axillae meristem organization and lateral organs. The findings reported herein may be useful for knowledge-based branch number of soybean improvement. Future research will focus on validating the effects of these putative major QTLs detected in this study. Our results demonstrated that comprehensive genetic architecture of branch number and characterizing the underlying candidate genes are powerful in simultaneous improvement and genetic dissection of complex traits such as branch number in soybean.

## Supporting information

S1 Table. List of consensus-QTLs after the integration of reproducible identified-QTLs for branch number.

S2 Table. Branch number QTLs of soybean in different studies

## Acknowledgments

This research was supported by the National Key Research and Development Program of China (2016YFD0101500, 2016YFD0101504) (http://www.most.gov.cn/) to RJZ, the funder had roles in study design, data collection and analysis, and preparation of the manuscript. This research was supported by Shanxi Provincial Natural Science Youth Fund (201901D211563) (http://kjt.shanxi.gov.cn/) to YHY, the funder had roles in study design, data collection and analysis, decision to publish and preparation of the manuscript. This research was supported by Shanxi Academy of Agricultural Science Postdoctoral Fund (YCX2020BH4) (http://www.sxagri.ac.cn/), the funder had a role in data collection and analysis, decision to publish and preparation of the manuscript. This research was supported by Shanxi Academy of Agricultural Science Doctoral Research Fund (YBSJJ1706) (http://www.sxagri.ac.cn/) to YHY, the funder had no role in study design, data collection and analysis, decision to publish, or preparation of the manuscript. This research was supported by Shanxi Academy of Agricultural Science Biological Breeding Engineering (17YZGC102) (http://www.sxagri.ac.cn/) to RJZ, the funder had no role in study design, data collection and analysis.

## Author Contributions

Conceived and designed the experiments: RJZ YHY YL. Performed the experiments: YHY YL. Analyzed the data: YHY YL. Contributed reagents/materials/analysis tools: YHY YL ZYB RJZ YCW. Wrote the paper: YHY YL.

